# Immunotherapy with nebulized pattern recognition receptor agonists restores severe immune paralysis and improves outcomes in mice with influenza-associated pulmonary aspergillosis

**DOI:** 10.1101/2024.10.02.616293

**Authors:** Jezreel Pantaleón García, Sebastian Wurster, Nathaniel D. Albert, Uddalak Bharadwaj, Keerthi Bhoda, Vikram K Kulkarni, Mbaya Ntita, Paris Rodríguez Carstens, Madeleine Burch-Eapen, Daniela Covarrubias López, Yongxing Wang, Dimitrios P. Kontoyiannis, Scott E. Evans

**Author notes:** Co-Corresponding authors: **Scott E. Evans, MD**, The University of Texas MD Anderson Cancer Center, Department of Pulmonary Medicine 6565 MD Anderson Blvd, Unit 1059, Houston, TX, 77030, United States of America., **Dimitrios P. Kontoyiannis, MD, ScD, PhD (Hon),** The University of Texas M.D. Anderson Cancer Center, Department of Infectious Diseases, Infection Control and Employee Health, 1515 Holcombe Boulevard, Unit 1460, Houston, TX, 77030, United States of America. Co-first authors (Jezreel Pantaleón García, Sebastian Wurster). Co-senior authors (Dimitrios P. Kontoyiannis, Scott E. Evans).

## Abstract

Influenza-associated pulmonary aspergillosis (IAPA) is a potentially deadly super-infection in patients with influenza pneumonia, especially those with severe disease, underlying immunosuppression, corticosteroid therapy, or requiring intensive care support. Given the high mortality of IAPA, adjunct immunomodulatory strategies remain a critical unmet need. Previously, desensitization of pattern recognition pathways has been described as a hallmark of IAPA pathogenesis and predictor of mortality in IAPA patients. Therefore, we studied the impact of nebulized Toll-like receptor 2/6/9 agonists Pam2 CSK4 (Pam2) and CpG oligodeoxynucleotides (ODN) on infection outcomes and pulmonary immunopathology in a corticosteroid-immunosuppressed murine IAPA model. Mice with IAPA receiving mock therapy showed rapidly progressing disease and a paralyzed immune response to secondary *A. fumigatus* infection. Nebulized Pam2ODN was well tolerated and significantly prolonged event-free survival. Specifically, dual-dose Pam2ODN therapy before and after *A. fumigatus* infection led to 81% survival and full recovery of all survivors. Additionally, transcriptional analysis of lung tissue homogenates revealed induction of PRR signaling and several key effector cytokine pathways after Pam2ODN therapy. Moreover, transcriptional and flow cytometric analyses suggested enhanced recruitment of macrophages, natural killer cells, and T cells in Pam2ODN-treated mice. Collectively, immunomodulatory treatment with nebulized Pam2ODN strongly improved morbidity and mortality outcomes and alleviated paralyzed antifungal immunity in an otherwise lethal IAPA model. These findings suggest that Pam2ODN might be a promising candidate for locally delivered immunomodulatory therapy to improve outcomes of virus-associated mold infections such as IAPA.

## Introduction

Lower respiratory tract infections (LRTIs) with respiratory viruses such as influenza A virus (IAV) are associated with significant morbidity and mortality, especially in immunocompromised hosts (1). Compounding the poor outcomes of viral LRTIs, secondary pneumonias caused by bacteria and, as increasingly appreciated, by fungi, constitute a significant source of morbidity and mortality after viral LRTIs (2, 3). Invasive pulmonary aspergillosis, predominantly caused by *Aspergillus fumigatus* (AF), is the commonest fungal super-infection after viral pneumonia (3). Concerningly, about 20% of critically ill influenza patients develop influenza-associated pulmonary aspergillosis (IAPA), and the incidence is even higher in patients with additional underlying risk factors for invasive aspergillosis and those receiving glucocorticosteroids for management of respiratory failure (4, 5). Even with modern intensive care and potent antifungal agents, mortality rates of IAPA remain as high as 50%, underscoring an unmet need for novel adjunct therapeutic approaches, including immunotherapy (5, 6).

The immunopathogenesis of virus-associated pulmonary aspergilloses is thought to be driven by a coalescence of virus-induced damage to the epithelial barrier, pulmonary hyperinflammation, and several hallmarks of local and systemic immune dysregulation and impairment (7–11). Common surrogates of pulmonary immune paralysis in IAPA patients and murine IAPA models include impaired recruitment and maturation of innate effector cells, unfavorable polarization and exhaustion of adaptive cellular immunity, and attenuated pattern recognition receptor (PRR) signaling (12–15). For instance, IAV infection reduced transcription levels of several PRRs in a murine IAPA model, including Toll-like receptors (TLRs) and C-type lectin receptors involved in fungal recognition (13). Likewise, transcriptional immune profiling of bronchoalveolar lavage (BAL) samples from IAPA patients revealed downregulation of several genes encoding proteins involved in fungal recognition and killing, including TLR2, a key receptor for recognition of AF conidia (14). Moreover, a recent study in IAPA patients admitted to the intensive care unit identified downregulation of PRR genes as a significant predictor of increased mortality (15), further corroborating the significance of impaired PRR signaling in the pathogenesis of IAPA. Therefore, immunomodulatory strategies restoring and enhancing pulmonary PRR signaling might be a promising adjunct approach to improve IAPA outcomes.

Prior studies by our group revealed significantly improved morbidity and mortality, enhanced epithelial resistance and viral clearance, and attenuated viral immune injury after inhaled immunomodulatory therapy with the synergistic TLR2/TLR6 and TLR9 agonists Pam2CSK4 and CpG oligodeoxynucleotides M362 (Pam2ODN) in murine models of various pneumonias, including IAV, paramyxovirus, and coronavirus infections (16–19). Furthermore, Pam2ODN provided a potent therapeutic benefit and facilitated rapid fungal killing in mice with underlying chemotherapy-induced immunosuppression and invasive pulmonary aspergillosis (20). Given these encouraging results in murine mono-infection models and the critical role of PRR signaling in the pathogenesis of IAPA, we herein studied Pam2ODN as an adjunct immunotherapy in mice with IAPA. We found that immunomodulatory treatment with nebulized Pam2ODN strongly improved infection outcomes and enhanced antimicrobial defense in our otherwise lethal and severely immuno-paralyzed corticosteroid-immunosuppressed IAPA model.

## Materials & Methods

### Murine infection model

Eight-to ten-week-old female BALB/c mice were infected with a sublethal dose (< LD_10_, ∼25,000 plaque forming units) of a mouse-adapted influenza A/Hong Kong/1968 (H3N2) strain, delivered by aerosolization, as previously described (21, 22). On days 5 and 8 after influenza infection, mice received two intra-peritoneal injections of 10 mg cortisone acetate (CA; Sigma-Aldrich). On day 9, mice were intranasally challenged with 50,000 AF-293 conidia or sterile saline (mock infection). For therapeutic experiments, mice received a 30-minute nebulization of either phosphate-buffered saline (PBS, mock treatment) or Pam2ODN (4 µM Pam2CSK4 + 1 µM ODN M362) before (single-dose, day 8) or before and after AF infection (dual-dose, days 8 and 12). Infection severity was scored daily using the viral pneumonia score (VPS, 0=healthy to 12=moribund), as previously described (23). To assess therapeutic efficacy, a combined morbidity and mortality endpoint was used, with an event defined as either death or reaching a VPS ≥7, a score indicating considerable distress and a high likelihood of death within 72 hours in the absence of anti-infective therapy (21, 23). This approach allowed us to obtain immunologic samples from non-moribund mice without compromising the primary endpoint analysis.

### Analysis of bronchoalveolar lavage fluid (BALF)

Sampling of BALF was performed by intratracheal instillation and collection of 1 mL of sterile PBS twice using a 20G cannula. Cytocentrifugation was performed with 200 μL of lavage fluid at 300 g for 5 minutes using a Cytospin 4 (Thermo Fisher Scientific). BALF samples were imaged in an automated BZX-810 microscope at 40x. Cell counting was performed in 2-3 FOV from independent mice from each experimental group. Individual cells in contact with the image edge, blurred cells with poorly defined borders, and large or dense clumps of cells were excluded. Leukocyte cells were differentially counted as macrophages, monocytes, lymphocytes and granulocytes (neutrophils or eosinophils). Total red blood cell counts included both erythrocytes and echinocytes (Burr cells).

### Transcriptional analyses

Lungs were weighed and flash-frozen in 1 mL RNAprotect Tissue Reagent (Qiagen). For RNA isolation, 1 mL RLT Buffer (Qiagen) was added to the thawed lungs, and tissue was homogenized using a Mini Bead-Beater (Biospec Products) and 15 acid-washed 3-mm glass beads (Sigma-Aldrich). Total RNA was isolated from a volume of lung homogenate equivalent to 30 mg of tissue using the RNeasy Tissue Kit and RNase-Free DNase Set (both Qiagen) according to the manufacturer’s instructions. Yield and purity of RNA were determined using a NanoDrop One^C^ spectrophotometer (Thermo Fisher Scientific).

Expression of 785 immune-related genes was then assessed using the murine Host Response nCounter panel (Nanostring Technologies). Data was analyzed using the nSolver Analysis Software, with background thresholding to the median of negative controls and normalization to the panel’s 12 housekeeping genes (geometric mean). Gene identifiers and pairwise mean inter-group expression ratios were imported into Ingenuity Pathway Analysis (Qiagen). Core analysis was performed to study canonical pathway enrichment, considering genes with mean inter-group expression fold changes of >|1.25| for infection-induced differences and >|1.5| for therapeutic effects. Differential enrichment of canonical pathways was considered significant at an absolute z-score value ≥ 1 and a Benjamini-Hochberg adjusted p value < 0.05.

### Flow cytometry

Single-cell lung suspensions were prepared from disaggregated lungs as previously reported (18). Briefly, murine lungs were harvested and stored in sterile PBS on ice for immediate processing. Lungs were cut into ≤1 mm^3^ pieces using razor blades and digested with collagenase/DNAse I at 5 mg/mL for 45 minutes at 37 °C. The resulting single-cell suspensions were passed through a 70 µm filter and washed with PBS. After red blood cell lysis, single-cell suspensions were washed with PBS + 1% fetal bovine serum and stained for specific cell markers. Antibody panels are summarized in **Table S1**. Cells were fixed and subsequently acquired on a BD LSRFortessa X-20 (BD Biosciences) flow cytometer.

### Cytokine measurements (Luminex assay)

Lung tissue was homogenized by bead beating in 1 mL PBS, as described above. A 13-plex magnetic murine cytokine & chemokine panel (LXSAMSM-13) was performed according to the manufacturer’s instructions (R&D Systems) and analyzed using a Luminex 200 device (Luminex Corporation). The following analytes were included: CCL2, CCL3, CCL4, CXCL2, GM-CSF, IFN-γ, IL-2, IL-4, IL-6, IL-12 p70, IL-17A, IL-33, and TNF-α.

### Statistical analyses

Survival curves and event-free survival were compared using the Mantel-Cox log-rank test. Results of immunoassays were compared using Student’s two-sided t-test or Mann-Whitney U test for two-group comparisons and one-way ANOVA with Tukey’s post-test or Kruskal-Wallis test with Dunn’s post-test for four-group comparisons. Significance tests are specified in the figure legends. Significant findings are denoted by asterisks: * p < 0.05, ** p < 0.01, *** p < 0.001. Numeric values are provided for p-values between 0.05 and 0.1. Data were analyzed and visualized using Excel 365 (Microsoft Corporation) and Prism v9 (GraphPad Software).

## Results

### CA-immunosuppressed mice IAPA display a state of severe immune paralysis

IAV-infected and CA-immunosuppressed mice showed rapidly progressing disease upon AF super-infection (i.e., induction of IAPA) (21). Despite severe infection, RT-qPCR analysis of lung tissue from mice with IAPA showed minimal induction of genes associated with key antifungal effector responses such as type-1 T-helper cell (Th1) (*IFNG*) and Th17 (*IL17A*) responses, mononuclear inflammation (*IL6*), and recruitment of innate effector cells (e.g., *CCL2*), with relative expression levels of 0.82-1.32 in IAPA vs. IAV only groups (**Fig. 1A**).

**Figure 1:**
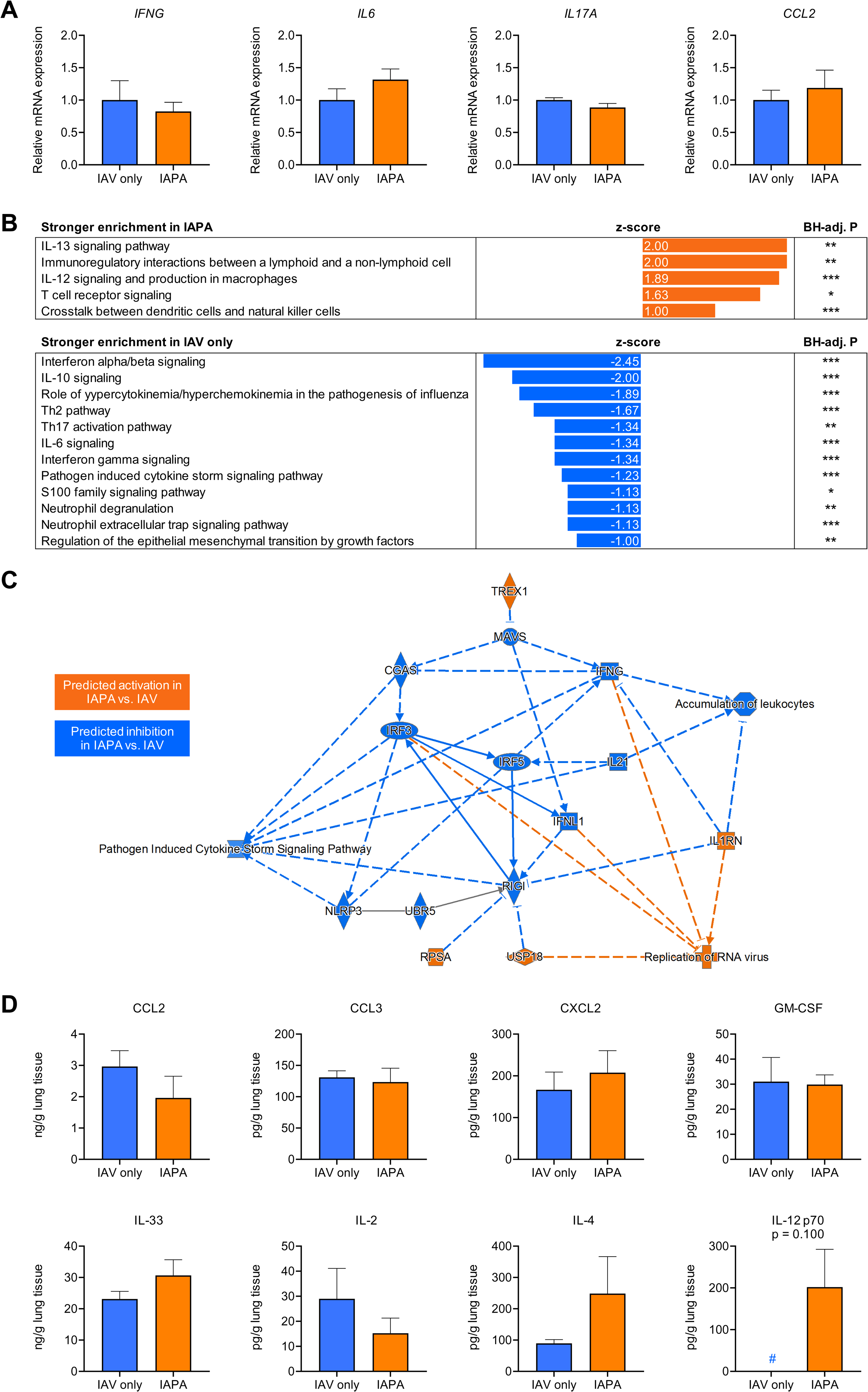
Mice with IAPA display an immuno-paralytic phenotype with dysregulated pattern recognition receptor signaling and minimal incremental inflammatory response to AF super-infection. (**A**) Expression of cytokine genes with known roles as master regulators of antifungal immunity in lung tissue homogenates of mice with IAPA compared to those with IAV mono-infection. Unpaired t-test. (**B**) Comparison of canonical pathway enrichment based on transcriptional responses (nCounter Host Response panel) in lung tissue of mice with IAPA compared to those with IAV mono-infection. Differential pathway enrichment was defined as an absolute z-score ≥ 1 and a Benjamini Hochberg (BH)-adjusted p-value ≤ 0.05. (**C**) Network of transcriptional changes to the pulmonary immune environment in mice with IAPA versus those with IAV mono-infection as predicted by Ingenuity Pathway Analysis. (**D**) Comparison of cytokine and chemokine concentrations in mice with IAPA versus those with IAV mono-infection. Mann-Whitney U test. # all 3 replicates in the “IAV only” group were below the lower limit of detection (∼ 30 pg/g lung tissue). TNF-α, IFN-γ, IL-6, IL-17A, and CCL4 were below the limit of detection in most or all mice, regardless of AF super-infection (not shown). (**A-D**) N = 3 mice per group and readout. Columns and error bars represent means and standard errors of the means, respectively. <u>Abbreviations:</u> AF = *Aspergillus fumigatus*, C(X)CL = C-(X-)C motif chemokine ligand, GM-CSF = granulocyte macrophage colony stimulating factor, IAPA = influenza-associated pulmonary aspergillosis, IAV = influenza A virus, IFN = interferon, IL = interleukin, Th = T-helper cell(s), TNF = tumor necrosis factor.

nCounter-based transcriptional analysis of lung tissue further corroborated a state of severe immune paralysis in mice with IAPA, with minimal inflammatory response to AF super-infection compared to IAV infection only. Specifically, only modest induction (Z-score, 1-2) of intercellular signaling and adaptive immune activation were seen upon AF super-infection (**Fig. 1B**). These signals were counterbalanced by significant suppression of several immune pathways in mice with IAPA versus those with IAV infection only, especially key effector cytokine pathways (e.g., IL-6, IFN-γ, type-1 interferons) and pathways associated with neutrophilic effector responses (**Fig. 1B**). Moreover, AF super-infection was associated with suppression of several mediators of PRR signaling pathways, including pathways associated with viral control (e.g., RIG-I, CGAS, IRF3, and IRF5) (**Fig. 1C**). These findings suggest a state of severe immune paralysis and impaired pathogen control in mice with IAPA.

Consistent with this observation, phenotypic validation of cytokine concentrations in lung tissue homogenates revealed largely comparable concentrations of innate effector cytokines in mice with IAPA versus those with IAV infection only. Specifically, no increased production of key chemokines and growth factors associated with recruitment of innate effector cells (CCL2, CCL3, CXCL2, GM-CSF and IL-33) was seen, with mean inter-group ratios of 0.66 – 1.33 (p = 0.400 – 1.000) (**Fig. 1D**). Moreover, type-1 and type-17 T-helper cell signature cytokines IFN-γ and IL-17A were below the detectable range in most animals, regardless of AF super-infection (data not shown), and pulmonary IL-2 concentrations tended to be lower in mice with IAPA compared to those with IAV infection only (**Fig. 1D**). Consistent with our nCounter-based pathway enrichment analysis, IL-12 p70 was the only tested cytokine markedly elevated after AF super-infection (**Fig. 1D**). Collectively, these findings corroborate a state of severe immune paralysis, with minimal incremental inflammation elicited by AF super-infection despite its severe “clinical” manifestation in our model.

### Immunotherapy with nebulized Pam2ODN improves clinical outcomes in immunosuppressed mice with IAPA

To overcome immune paralysis in mice with IAPA and re-invigorate PRR signaling, mice received nebulized immunostimulatory Pam2ODN treatment either before AF super-infection or both before and after super-infection (**Fig. 2A**). All mock-treated corticosteroid-immunosuppressed mice with IAPA reached the combined morbidity/mortality endpoint by day 13, i.e., within 4 days of AF infection. Single-dose Pam2ODN therapy on day 8 led to universal event-free survival until day 13 but all mice reached the morbidity/mortality endpoint by day 16 (**Fig. 2B**). In contrast, dual-dose Pam2ODN therapy on days 8 and 12, i.e., both before and after *A. fumigatus* infection, led to 80% event-free survival until day 21 (**Fig. 2B**, p < 0.001 versus all other groups). Notably, all survivors fully recovered by day 21 (mean VPS = 0.4, **Fig. 2C**). Consistent with these clinical trends, microscopic analysis of BAL fluid corroborated markedly reduced erythrocyte counts as a surrogate of hemorrhagic lesions in Pam2ODN-treated animals with IAPA (means, 43 and 18 after single-dose and dual-dose Pam2ODN, respectively) compared to mock-treated infected animals (mean, 275; **Fig. 2D**).

**Figure 2:**
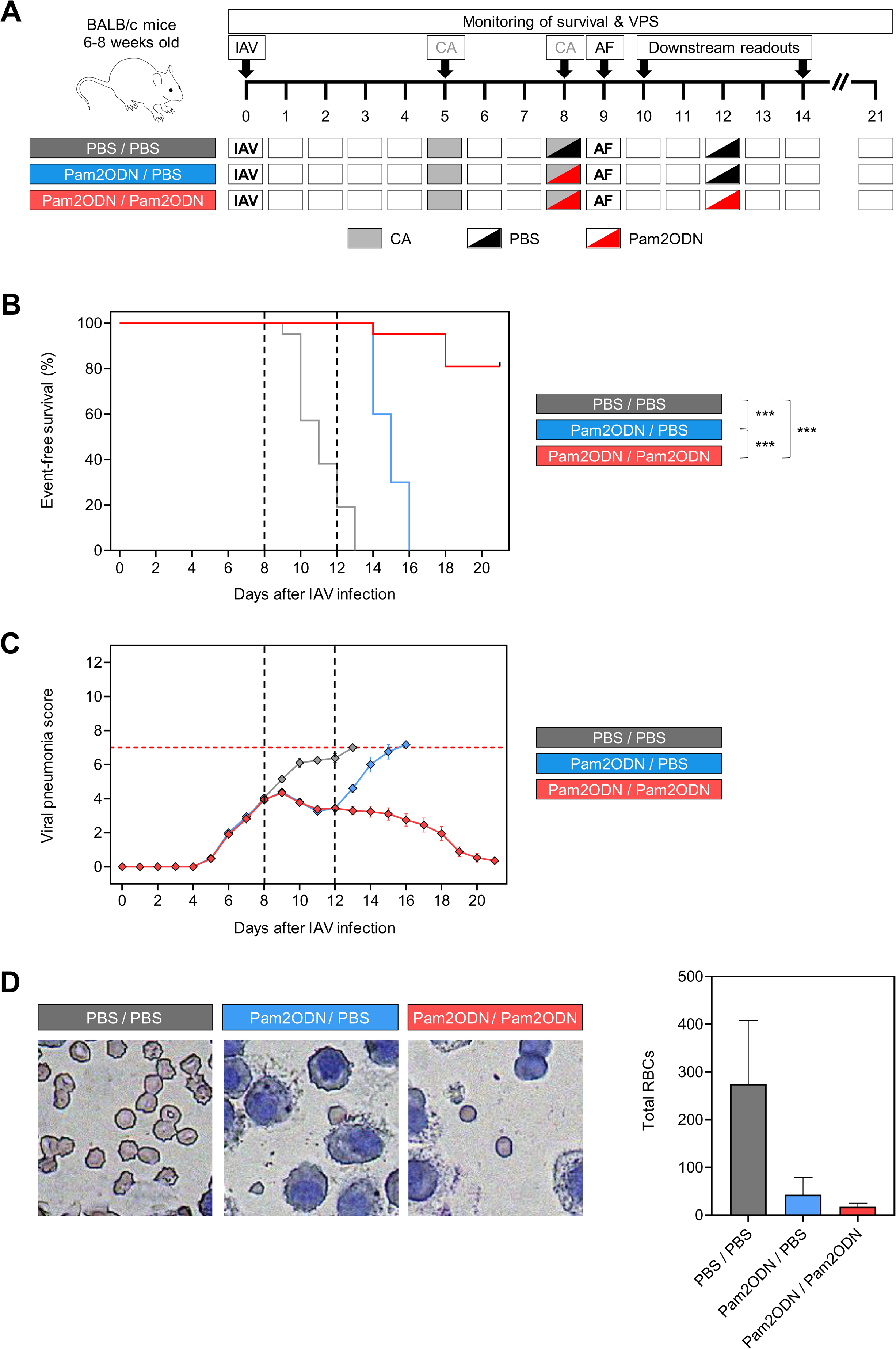
Dual-dose immunotherapy with nebulized Pam2ODN strongly improves clinical outcomes in CA-immunosuppressed mice with IAPA. (**A**) Outline of experimental interventions. (**B**) Kaplan-Meier curves comparing event-free survival in CA-immunosuppressed mice with IAPA according to the treatment arm. N =20-21 mice per group assessed across 3 independent experiments. Mantel-Cox log-rank test. (**C**) VPS kinetics according to the treatment arm. Mice that have reached the morbidity/mortality endpoint prior to the time of assessment are excluded. (**D**) Representative micrographs of RBC burden in bronchoalveolar lavage and quantitative analysis of total RBCs per field according to the treatment arm on day 14 (5 days after AF infection). N = 3 mice per treatment arm. (**C-D**) Means and standard errors of the mean (error bars) are shown. <u>Abbreviations:</u> AF = *Aspergillus fumigatus*, CA = cortisone acetate, IAPA = influenza-associated pulmonary aspergillosis, IAV = influenza A virus, PBS = phosphate-buffered saline, Pam2ODN = Pam-2 CSK4 + CpG oligodeoxynucleotides M362, RBC(s) = red blood cell(s), VPS = viral pneumonia score.

### Immunotherapy with Pam2ODN re-invigorates PRR and downstream cytokine signaling

To comprehensively characterize the pulmonary immune environment in mock- and Pam2ODN-treated mice with IAPA, we performed nCounter and pathway enrichment analysis. Dual-dose Pam2ODN therapy led to significant induction of PRR (TLR, NOD1/2, cGAS/STING) and NF-κB signaling pathways, along with induction of key innate effector cytokine pathways (e.g., IL-6, IL-8, IL-17A, IL-33) (**Fig. 3A**). These changes were associated with several signals of enhanced innate immune cell effector responses, including enhanced DC maturation, macrophage activation, phagosome formation, and oxidative burst, as well enhanced neutrophil degranulation and neutrophil extracellular trap signaling (**Fig. 3A**). Furthermore, pathway enrichment analysis suggested modest enhancement of T-helper signaling (**Fig. 3A**). Notably, 7 out of the 12 pathways significantly suppressed in mice with IAPA compared to those with IAV mono-infection (**Fig. 1B**) were restored by dual-dose Pam2ODN therapy (**Fig. 3A**).

**Figure 3:**
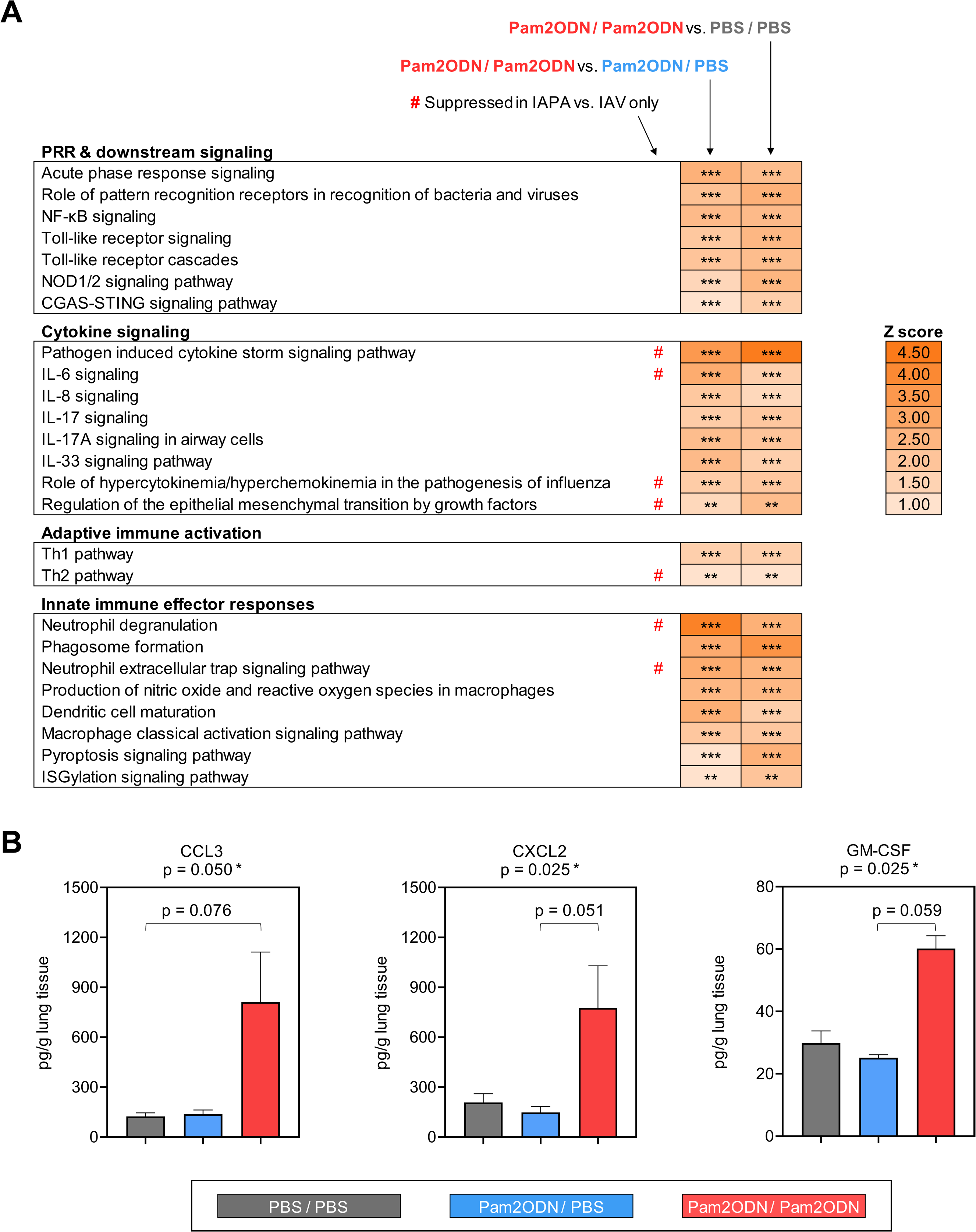
Dual-dose immunotherapy with nebulized Pam2ODN before and after AF super-infection enhances innate immune defense in CA-immunosuppressed mice with IAPA. (**A**) Comparison of canonical pathway enrichment based on transcriptional responses (nCounter Host Response panel) in lung tissue homogenates of mice with IAPA according to the treatment arm. (**B**) Concentrations of selected cytokines/chemokines in lung tissue homogenates of mice with IAPA according to the treatment arm. Kruskal-Wallis test with Dunn’s post-test. All other tested cytokines were either below the limit of detection in most mice regardless of the treatment arm (CCL4, IFN-γ, IL-6, IL-12 p70, IL-17A, and TNF-α) or not significantly different between the treatment arms (CCL2, IL-2, IL-4, and IL-33) (**Table S2**). (**A-B**) All analyses were performed on day 14 (5 days after AF infection). N = 3 mice per group and readout. <u>Abbreviations:</u> AF = *Aspergillus fumigatus*, CGAS-STING pathway = cyclic GMP-AMP synthase / stimulator of interferon genes pathway, C(X)CL = C-(X-)C motif chemokine ligand, GM-CSF = granulocyte macrophage colony stimulating factor, IAPA = influenza-associated pulmonary aspergillosis, IAV = influenza A virus, IFN = interferon, IL = interleukin, NF-κB = nuclear factor kappa-light-chain-enhancer of activated B-cells, NOD1/2 = nucleotide-binding oligomerization domain-containing protein 1/2, PBS = phosphate-buffered saline, Pam2ODN = Pam-2 CSK4 + CpG oligodeoxynucleotides M362, Th = T-helper cell(s).

An integrative predicted network of gene- and pathway-level expression changes corroborated enhanced PRR and downstream signaling (e.g., MYD88, RelA) after dual-dose Pam2ODN treatment. Furthermore, Ingenuity Pathway Analysis revealed induction of several key effector cytokines (e.g., TNF-α, IFN-γ, IL-1β, IL-6) and chemokines (CXCL2, CXCL3) that are associated with antifungal immune enhancement, mobilization of innate effector cells, and increased pathogen clearance (**Fig. S1**) (24).

### Pam2ODN promotes accumulation of mature mononuclear phagocytes, natural killer cells, and T cells in the lung

Next, we tested induction of antifungal effector cytokines on protein level by Luminex-based analysis of lung tissue homogenates. While less sensitive than our transcriptional analyses, we found strong induction of cytokines associated with mobilization of innate effector cells, especially mononuclear phagocytes, in mice with IAPA that received dual-dose Pam2ODN therapy compared to those receiving the mock treatment or single-dose Pam2ODN (**Table S2**). Specifically, strong and significant elevations of CCL3 (6.57-fold compared to mock therapy, p = 0.025), CXCL2 (3.74-fold, p = 0.050), and GM-CSF (2.01-fold, p = 0.025) were found after dual-dose Pam2ODN therapy (**Fig. 3B**). Consistent with this observation, Ingenuity Pathway Analysis of transcriptional data predicted increased mobilization of mononuclear phagocytes, natural killer cells, and (T) lymphocytes after dual-dose Pam2ODN compared to mock therapy and single-dose Pam2ODN treatment (**Fig. S1**).

Given the strong impact of the second (post-AF) Pam2ODN dose on pulmonary secretion of cytokines and chemokines associated with recruitment of innate effector cells, we used flow cytometry to compare the cellular composition of lung tissue from mice that received dual-dose vs. single-dose Pam2ODN therapy. Compared to single-dose therapy, dual-dose Pam2ODN led to globally increased leukocyte-to-epithelial cell ratios in lung tissue homogenates (1.79 vs. 1.08, **Fig. 4A**). Particularly, mice with IAPA that received dual-dose Pam2ODN therapy showed markedly enhanced pulmonary recruitment of alveolar macrophages (6.2% vs. 1.6% of total viable cells, p = 0.042), interstitial macrophages (5.3% vs. 2.4%, p = 0.031), CD11b^-^ dendritic cells (1.5% vs. 0.3%, p = 0.018), natural killer cells (4.2% vs. 1.5%, p = 0.056), and T-lymphocytes (10.9% vs. 4.7%, p = 0.011) compared to mice that received single-dose Pam2ODN (**Fig. 4B**). Of note, due to the low number of non-moribund animals, flow cytometric data from the mock-treated group (PBS/PBS) could not be obtained.

**Figure 4:**
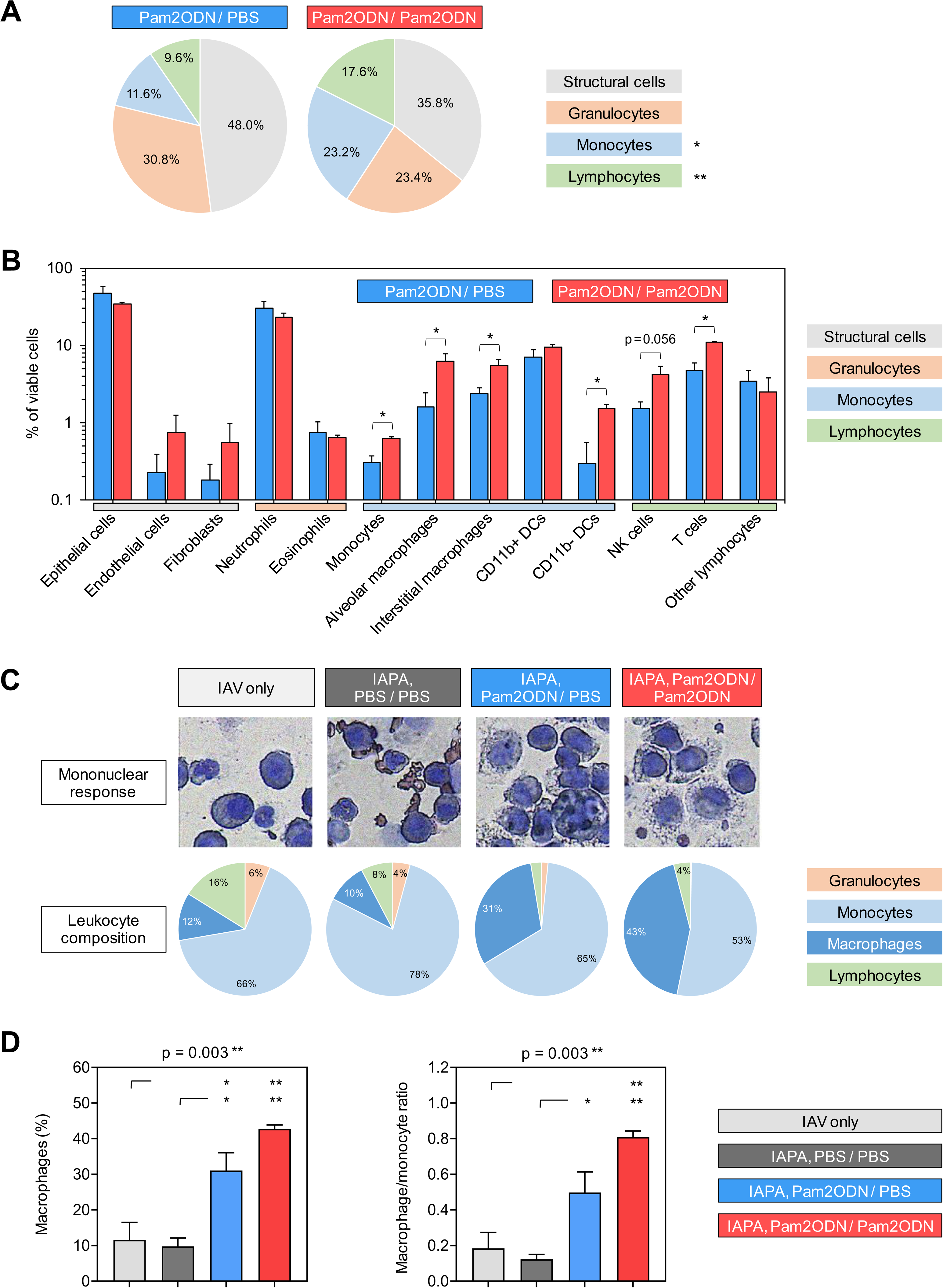
Immunotherapy with nebulized Pam2ODN promotes pulmonary recruitment of mature mononuclear effector cells in mice with IAPA. (**A-B**) Flow cytometric analysis of immune cells and structural cell populations (epithelium, endothelium, fibroblasts) in lung tissue homogenates of mice with IAPA receiving single-dose versus dual-dose Pam2ODN therapy. Unpaired t-test. All significant results were confirmed to have false discovery rates < 0.2 (Benjamini-Hochberg method). (**C**) Representative images and quantitative analysis of leukocyte subsets in BAL according to the infection and treatment arm. Subsets ≤ 3% are not labelled. **(D)** Proportions of macrophages among leukocytes and macrophage/monocyte ratios in BAL according to the infection and treatment arm. One-way analysis of variance with Tukey’s post-test. (**A-D**) N = 3 mice per treatment arm and assay. All analyses were performed on day 14 (5 days after AF infection). <u>Abbreviations:</u> AF = *Aspergillus fumigatus*, BAL = bronchoalveolar lavage, CD = cluster of differentiation, DCs = dendritic cells, IAPA = influenza-associated pulmonary aspergillosis, IAV = influenza A virus, NK cells = natural killer cells, PBS = phosphate-buffered saline, Pam2ODN = Pam-2 CSK4 + CpG oligodeoxynucleotides M362.

To further validate Pam2ODN-induced changes to the pulmonary leukocyte repertoire, we performed microscopic analysis of leukocyte composition in BALF. Here, we found a significant increase in macrophages from 10% in mock-treated mice with IAPA to 31% and 43% in those receiving single- and dual-dose Pam2ODN therapy, respectively (p=0.003, **Fig. 4C-D**). This was paralleled by an increase in the macrophage/monocyte ratio from 0.13 (mock treatment) to 0.48 (single-dose Pam2ODN) and 0.81 (dual-dose Pam2ODN), respectively (p=0.003, **Fig. 4E**). These findings corroborate that immune protection by Pam2ODN therapy is, at least in part, driven by increased mobilization and maturation of mononuclear phagocytes.

Altogether, our data support a model whereby immunotherapy with nebulized Pam2ODN re-invigorates PRR signaling, enhances mobilization of key immune cell populations, and induces innate effector cytokine responses (**Fig. 5**). In combination with the known role of Pam2ODN as an inducer of epithelial resilience, these changes to the pulmonary immune environment alleviate infection-induced immune paralysis and restore anti-AF defense in mice with IAPA, thereby strongly improving morbidity/mortality outcomes.

**Figure 5:**
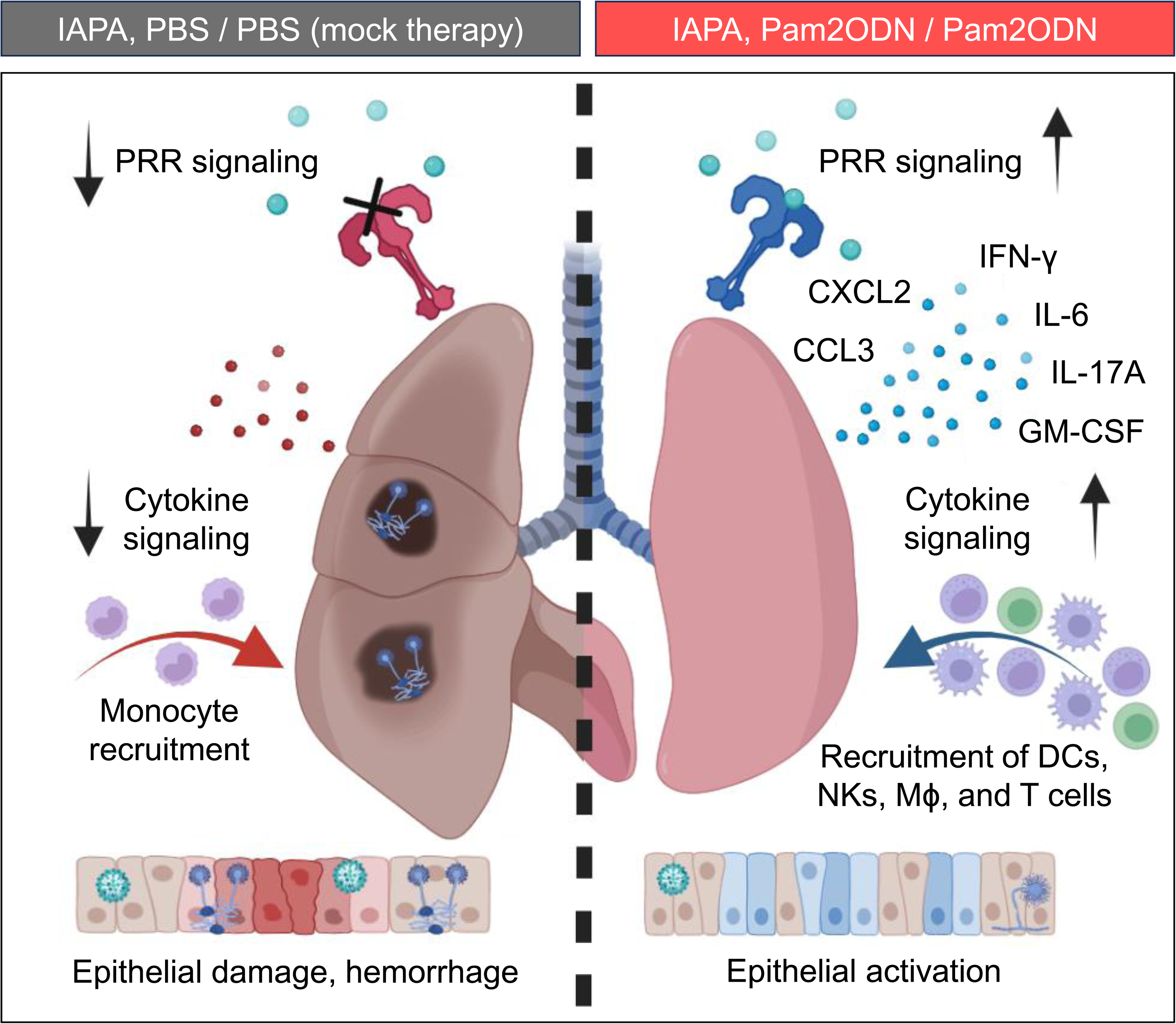
Immunotherapy with nebulized Pam2ODN alleviates pulmonary immunopathology to improve outcomes in immunosuppressed mice with IAPA. Schematic summarizing immuno-protective effects of nebulized Pam2ODN therapy in mice with IAPA. Abbreviations: C(X)CL = C-(X-)C motif chemokine ligand, GM-CSF = Granulocyte-macrophage colony-stimulating factor, IAPA = influenza-associated pulmonary aspergillosis, IFN = interferon, IL = interleukin, Mϕ = macrophages, NK = natural killer cells, PBS = phosphate-buffered saline, Pam2ODN = Pam-2 CSK4 + CpG oligodeoxynucleotides M362, PRR = Pattern recognition receptor.

## Discussion

The COVID-19 pandemic has highlighted the significant healthcare burden, morbidity, and mortality caused by secondary fungal pneumonias. Given the heightened prevalence of respiratory viruses associated with severe secondary mold pneumonias, novel adjunct strategies are needed to improve the outcomes of these infections. Here, we studied locally delivered (nebulized) PRR agonists as an immunotherapeutic intervention to improve the outcome of IAPA in glucocorticosteroid-immunosuppressed mice by attenuating epithelial pathogenesis and pulmonary immune paralysis.

In the first part of this study, we characterized the natural response to AF super-infection in mice with underlying influenza and high-dose corticosteroid therapy. Our observation of a paralyzed or even suppressed immune environment in mice with IAPA *versus* those with influenza only aligns with extensive evidence from immune monitoring studies in human patients with viral pneumonia and post-viral mold infections. For instance, we found a notable overlap of the most strongly suppressed transcriptional pathways (e.g., type-1 interferon signaling, IFN-γ signaling, IL-6 signaling, and various pathways related to neutrophil effector responses) between our model and a prior study in human patients with IAPA *versus* influenza only using a similar methodology (nCounter-based transcriptomics with pathway enrichment analysis) (14). Our finding of a largely anergic immune state and impaired PRR sensing is also consistent with a more recent publication by the same group showing minimal inflammatory responses to bacterial co-infection in influenza patients with and without IAPA (25). Additionally, our finding of impaired viral pathogen sensing via RIG-I, cGAS, and their downstream effectors in mice with IAPA *versus* those with influenza mono-infection aligns well with the mutual impairment of viral and fungal PRR pathways previously described in an *in-vitro* model of AF and cytomegalovirus co-infection (26).

Our model used CA immunosuppression to recapitulate a common risk factor in human IAPA patients and facilitate robust infection with a relatively low AF inoculum. While it is conceivable that CA contributed to the lack of a pronounced inflammatory response to AF super-infection in our model, there are several lines of evidence suggesting that CA was not a major confounder of our findings. On the one hand, potent innate immune cell recruitment and cytokine release had been reported previously in single-infection pulmonary aspergillosis models using similar high-dose glucocorticosteroid regimens (27, 28). On the other hand, several clinical immune phenotyping studies suggested that hallmarks of paralyzed antifungal immunity in patients with viral pneumonia were largely independent of corticosteroid therapy (14, 29). Furthermore, our findings of both attenuated PRR signaling and transcriptional surrogates of impaired neutrophil effector responses in mice with IAPA align with data from Liu and colleagues who used an IAPA model without pharmacological immunosuppression (13). Similarly, Lee and colleagues found that inflammation in a non-immunosuppressed IAPA model was mainly driven by the underlying IAV infection, whereas secondary aspergillosis had a limited contribution to inflammatory responses (30).

To overcome dysfunctional PRR signaling and immune paralysis in our CA-immunosuppressed IAPA model, we tested immunomodulatory therapy with nebulized Pam2ODN, which allowed most mice to clear IAPA infection and fully recover from infection-induced distress after only two Pam2ODN doses and without concomitant antiviral or antifungal therapy. Mechanistically, dual-dose Pam2ODN immunotherapy enhanced PRR and downstream signaling, induced essential antifungal effector cytokine pathways, and promoted recruitment, differentiation, and maturation of mononuclear phagocytes. Thereby, Pam2ODN immunotherapy targets multiple common immune deficits in sequential inter-kingdom infections (7–9).

Interestingly, single-dose Pam2ODN before AF super-infection elicited minimal changes to pro-inflammatory cytokine release. This might be due to the timing of the first Pam2ODN dose after the peak of IAV-induced distress and before AF super-infection. Additionally, given that PRR agonists are known inducers of trained immunity (31), the leukocyte-driven component of Pam2ODN-induced immune augmentation likely benefits from repeat exposure to the agent, as seen previously in mice treated with individual nebulized exposures to Pam2, ODN, and other PRR agonists at increasing concentrations (16).

While the present study focused on leukocyte-driven pulmonary immune augmentation, Pam2ODN was previously shown to induce epithelial resistance in various preclinical studies of bacterial and viral mono-infections, partially due to its capacity to stimulate mitochondrial production of reactive oxygen species (ROS) and induce ROS-dependent epithelial STAT3 signaling (32, 33).

Consistent with the favorable tolerability data for nebulized Pam2ODN (PUL-042) in human patients with viral pneumonia (ClinicalTrials.gov NCT04312997) and our extensive prior preclinical work in various single-infection pneumonia models, immunotherapy was well-tolerated in our IAPA model and no evident immunotoxicities were seen. In fact, the conceivable tolerability advantages of topically delivered immunomodulators compared to systemic immune enhancers studied as antifungal immunotherapies might be particularly relevant in a background of underlying viral infections that are often associated with systemic hyperinflammation. Therefore, our favorable tolerability and efficacy data might encourage further comparative investigations of inhaled versus systemic administration of immunomodulators in primary or secondary mold pneumonias or polymicrobial respiratory tract infections. For instance, immune checkpoint inhibitors or recombinant GM-CSF showed promise as antifungal immunotherapies (34–37) and are available as (investigational) nebulized immune agents (38–39).

Furthermore, the benefits of Pam2ODN might also apply to other co- and sequential infections. Feys and colleagues found that several signals of immune paralysis, including suppressed TLR2 expression, were shared between patients with IAPA and COVID-19-associated pulmonary aspergillosis (14). Similarly, as discussed above, strong mutual impairment of PRR signaling and anti-infective effector responses was found in an *in-vitro* model of *A. fumigatus* and cytomegalovirus co-infection (26). Additionally, post-viral impairment of antimicrobial signaling has also been described in response to other fungal stimuli. For instance, weak PRR expression and dysfunctional phagocytosis after IAV infection were encountered in mice challenged with either *A. fumigatus* conidia or yeast zymosan (13). Likewise, patients with moderate COVID-19 showed surrogates of impaired TLR signaling in response to *Rhizopus arrhizus*, the commonest cause of COVID-19-associated mucormycosis (11, 29). Collectively, these studies suggest that dysfunctional PRR signaling may be a hallmark of immune dysregulation that is broadly applicable to many viral and fungal inter-kingdom infection settings, inviting further studies of Pam2ODN immunotherapy in other co- and sequential infection models, e.g., highly lethal mold pneumonias after LRTIs due to respiratory syncytial virus (40). Moreover, patients with IAPA and bacterial co-infection were shown to have minimal incremental pro-inflammatory pulmonary cytokine responses to bacterial co-pathogens, corroborating a severely paralyzed immune state (25). As Pam2ODN confers robust protection against virulent gram-positive and gram-negative bacteria (16), nebulized Pam2ODN might be a particularly attractive immunotherapeutic approach in patients with complex polymicrobial pneumonias due to its potential triple activity against viral, fungal, and bacterial co-pathogens.

This proof-of-concept study has several limitations. Immune impairment and immunomodulatory therapy were studied without the complexities of conventional antiviral or antifungal agents that would modulate infection severity and inflammation but might also elicit both immunostimulatory and inhibitory effects (21, 41–43). Furthermore, our study utilized a single IAV and AF strain and therefore does not reflect the considerable strain-to-strain variability. Despite the consistency of the underlying immune paralysis phenotype in our IAPA model with both previously published mouse models using different IAV and AF strains and real-life human patient data, confirmatory evidence for the benefits of Pam2ODN therapy against sequential infection with additional strains would be warranted. Moreover, the high mortality and rapid disease progression in the mock-treated IAPA group greatly limited sample availability for our various downstream readouts, especially on day 14 (day 5 after AF super-infection), reducing statistical power and not allowing us to perform all assays on the mock-treated cohort. Lastly, secondary mold pneumonias after respiratory viral infections most commonly affect patients with underlying classical risk factors for invasive fungal infections, such as uncontrolled diabetes mellitus, hematological malignancies, transplant history, or ongoing immunosuppressive therapies. Besides corticosteroid therapy, such comorbidities were not recapitulated in our study using *a priori* healthy inbred mice.

Collectively, our findings corroborate that IAPA is associated with a severely immuno-paralytical state, aligning with prior studies in IAPA mouse models and human patients with post-viral mold pneumonia. Immunotherapy with nebulized Pam2ODN was well tolerated and strongly improved clinical outcomes in our otherwise highly lethal IAPA model, enhanced pulmonary recruitment of mature mononuclear phagocytes, and induced antimicrobial signaling. These findings suggest a promising therapeutic potential of nebulized immunomodulators to mitigate pulmonary immune paralysis and improve IAPA outcomes.

## Acknowledgments

This study was partially supported by an investigator-initiated grant from Gilead Global Pharma (Grant IN-US-131-5756 to DPK and SW). Additional support was provided by the University Cancer Foundation via the Institutional Research Grant program at the University of Texas MD Anderson Cancer Center (to DPK), the Robert C. Hickey Chair for Clinical Care endowment (to DPK), and the Cyrus Scholar Award (to SW). This study was also supported by the National Institutes of Health (Grant R35 HL144805 to SEE).

## Meetings where the information has previously been presented

Preliminary data for parts of this study were presented at Trends in Medical Mycology 2023, Athens, Greece; the Biennial Symposium of the Mycoses Study Group Education and Research Consortium 2024, Colorado Springs, USA; and the American Thoracic Society International Conference 2023 (Washington, USA) and 2024 (San Diego, USA).

