## Supplementary material for "Immunotherapy with nebulized pattern recognition receptor agonists restores severe immune paralysis and improves outcomes in mice with influenza-associated pulmonary aspergillosis": S1

Supplementary Materials

**Table S1. Flow cytometry antibodies and reagents.**

| Materials / Resources | Reference/Identifier | Source |
|---|---|---|
| ***Conventional Flow Cytometry: Myeloid Antibody Panel*** | | |
| CD45 (30-F11) – redFluor 700 | Cat # 80-0451-U100 | Tonbo Biosciences |
| CD11b (M1/70) – PerCP-Cy5.5 | Cat # 45-0112-82 | Invitrogen |
| CD11c (N418) – Brilliant Violet 421 | Cat # 117330 | BioLegend |
| MHC II (M5/114.15.2) – PE-Cy 7 | Cat # 107630 | BioLegend |
| F4/80 Antigen (BM8.1) – APC-Cy 7 | Cat # 25-4801-U100 | Tonbo Biosciences |
| Ly-6 C (HK1.4) – APC | Cat # 128016 | BioLegend |
| Ly-6 G (1A8) – Alexa Fluor 488 | Cat # 127626 | BioLegend |
| CD24 (M1/69) – PE | Cat # 50-0242-U100 | Tonbo Biosciences |
| CD64 (X54-5/7.1) – Brilliant Violet 711 | Cat # 139311 | BioLegend |
| Siglec-F (E50-2440) – PE-CF594 | Cat # 562757 | BD Biosciences |
| Ghost Dye Violet 510 | Cat # 13-0870-T100 | Tonbo Biosciences |
| ***Conventional Flow Cytometry: T Cells Antibody Panel*** | | |
| CD3e (145-2C11) – PerCP-Cy 5.5 | Cat # 65-0031-U100 | Tonbo Biosciences |
| UCD4 (RM4-5) – PE-Cy 7 | Cat # 60-0042-U100 | Tonbo Biosciences |
| CD8a (53.6.7) – APC-Cy 7 | Cat # 25-0081-U100 | Tonbo Biosciences |
| CD44 (IM7) – Brilliant Violet 711 | Cat # 103057 | BioLegend |
| CD62L (MEL-14) – Brilliant Violet 421 | Cat # 104435 | BioLegend |
| CD25 (PC61) – Brilliant Violet 650 | Cat # 102037 | BioLegend |
| ROR γ T (AFKJS-9) – APC | Cat # 17-6988-82 | Invitrogen |
| FOXP3 (FJK-16s) – PE | Cat # 12-5773-82 | Invitrogen |
| Gata-3 (TWAJ) – Alexa Fluor 488 | Cat # 53-9966-42 | Invitrogen |
| Ghost Dye Violet 510 | Cat # 13-0870-T100 | Tonbo Biosciences |
| T-bet (4B10) – Brilliant Violet 785 | Cat # 644835 | BioLegend |
| CD45 (30-F11) – redFluor 700 | Cat # 80-0451-U100 | Tonbo Biosciences |
| ***Imaging Flow Cytometry: Structural Cells Antibody Panel*** | | |
| DAPI solution, 1mg/mL | Cat # 62248 | Thermo Fisher Scientific |
| CD45 (30-F11) – Brilliant Violet 605 | Cat # 103140 | BioLegend |
| CD324 (DECMA-1) – PE-Dazzle 594 | Cat # 147316 | BioLegend |
| Gp38 (8.1.1) – PerCP-Cy 5.5 | Cat # 127422 | BioLegend |
| CD31 (MEC13.3) – APC-Cy 7 | Cat # 102534 | BioLegend |
| NF-kB p65 (F-6) – Alexa Fluor 488 | Cat # sc-8008 AF488 | Santa Cruz Biotechnology |
| c-Jun (G-4) – Alexa Fluor 647 | Cat # sc-74543 AF647 | Santa Cruz Biotechnology |

Abbreviations: CD = Cluster of Differentiation; PerCP = Peridinin-Chlorophyll-Protein; MHC = Major Histocompatibility Complex; Ly-6 = Lymphocyte Antigen 6 Complex; Siglec = Sialic Acid Binding Ig-like Lectin; ROR = Retinoic Acid Receptor-related Orphan Receptor; FOXP3 = Forkhead Box P3; PE = Phycoerythrin; Cy = Cyanine; CF = Cyanine-based Fluorescent; APC = Allophycocyanin; T-bet = T-box expressed in T cells.

**Table S2. Cytokine concentrations in lung tissue of mice with IAPA according to the treatment arm.**

All concentrations, including the LOD, are provided as pg per g of lung tissue.

| Treatment ▶ | | PBS / PBS | Pam2ODN / PBS | Pam2ODN / Pam2ODN | | |
|---|---|---|---|---|---|---|
| **Analyte** | **LOD [a]** | **Mean** | **Mean** | **Mean** | **Ratio vs. PBS / PBS** | **Ratio vs. Pam2ODN / PBS** |
| CCL2 | 389.2 | 1962.5 | # (2019.7) | 2488.0 | 1.27 | # (1.23) |
| CCL3 | 1.0 | 123.4 | 138.0 | 811.0 | 6.57 | 5.88 |
| CCL4 | 181.5 | Below the LOD in most mice or in at least one mouse per group. | | | | |
| CXCL2 | 4.9 | 207.5 | 148.0 | 776.7 | 3.74 | 5.25 |
| GM-CSF | 10.3 | 29.9 | 25.1 | 60.1 | 2.01 | 2.39 |
| IFN-γ | 17.4 | Below the LOD in most mice or in at least one mouse per group. | | | | |
| IL-2 | 10.0 | 15.2 | 18.8 | 20.7 | 1.36 | 1.11 |
| IL-4 | 59.8 | 248.6 | 371.0 | 179.5 | 0.72 | 0.48 |
| IL-6 | 28.8 | Below the LOD in most mice or in at least one mouse per group. | | | | |
| IL-12 p70 | 46.8 | Below the LOD in most mice or in at least one mouse per group. | | | | |
| IL-17A | 45.2 | Below the LOD in most mice or in at least one mouse per group. | | | | |
| IL-33 | 103.9 | 30649.7 | 27699.2 | 38432.2 | 1.25 | 1.39 |
| TNF-α | 2.1 | Below the LOD in most mice or in at least one mouse per group. | | | | |

[a] 10% of the lowest standard at 100 mg lung weight.

#: Mean and ratio not reliably determinable due to one measurement below the LOD. Numbers in parentheses indicate the mean when excluding the single low outlier.

Abbreviations: C(X)CL = C-(X-)C motif chemokine ligand, GM-CSF = Granulocyte-macrophage colony-stimulating factor, IAPA = influenza-associated pulmonary aspergillosis, IFN = interferon, IL = interleukin, LOD = limit of detection, PBS = phosphate-buffered saline, Pam2ODN = Pam-2 CSK4 + CpG oligodeoxynucleotides M362, TNF-α = tumor necrosis factor alpha.

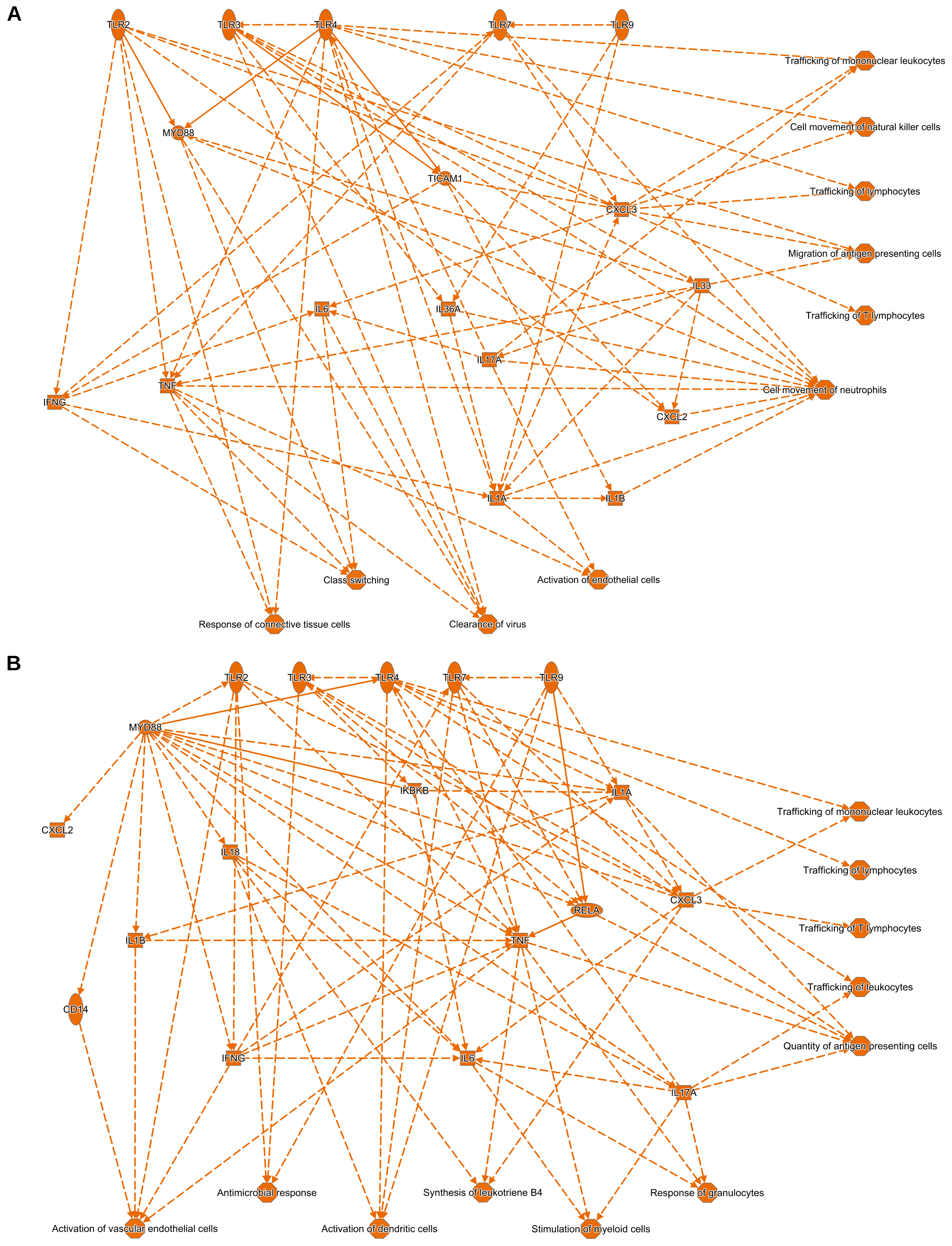


**Figure S1 (caption on following page)**

**Figure S1: Dual-dose Pam2ODN immunotherapy induces a transcriptional framework promoting recruitment of mononuclear effector cells to the lungs of mice with IAPA.**

Networks of transcriptional changes to the pulmonary immune environment in mice receiving dual-dose Pam2ODN therapy compared to those receiving either single-dose Pam2ODN (**A**) or mock therapy (**B**), as predicted by Ingenuity Pathway Analysis. N = 3 mice per treatment arm.

Abbreviations: C(X)CL = C-(X-)C motif chemokine ligand, IAPA = influenza-associated pulmonary aspergillosis, IFN = interferon, IKBKB = inhibitor of nuclear factor kappa B kinase subunit beta, IL = interleukin, MYD88 = myeloid differentiation primary response 88, NK = natural killer cells, PBS = phosphate-buffered saline, Pam2ODN = Pam-2 CSK4 + CpG oligodeoxynucleotides M362, RELA = v-rel avian reticuloendotheliosis viral oncogene homolog A, TICAM1 = TIR domain containing adaptor molecule 1, TLR = Toll-like receptor.
